# FAM134B isoform 2/RETREG1-2 defines a calnexin–TOLLIP–coupled ER-phagy pathway that restricts Ebola virus glycoprotein and is antagonized by VP40 through macro-autophagy

**DOI:** 10.64898/2026.04.01.715898

**Authors:** Jing Zhang, Tao Wang, Jiaxin Wen, Jing Lan, Sunan Li, Yong-Hui Zheng

**Affiliations:** State Key Laboratory of Animal Disease Control and Prevention, Harbin Veterinary Research Institute, Chinese Academy of Agricultural Sciences, Harbin, China; Department of Microbiology and Immunology, The University of Illinois Chicago, Chicago, IL 60612

**Author notes:** These authors contributed equally (Jing Zhang, Tao Wang).

**Keywords:** RETREG1, FAM134, TOLLIP, calnexin, Ebola, glycoprotein, ER-phagy

## Abstract

Selective autophagy of the endoplasmic reticulum (ER-phagy) is critical for ER proteostasis and host defense, yet how ER quality-control pathways interface with ER-phagy to restrict viral glycoproteins remains poorly defined. Previously, the 1^st^ known ER-phagy receptor gene RETREG1 (RETR1)/FAM134B gene was reported to restrict Ebola virus (EBOV) replication in vivo by inhibiting the viral glycoprotein (GP) and viral protein 40 kDa (VP40) expression, but this mechanism remains unknown. Here, we identify the truncated RETR1/FAM134B isoform 2 (RETR1-2), but not its full-length protein RETR1, as an ER-phagy receptor that targets EBOV-GP for degradation. RETR1-2 broadly triggers GP degradation across ebolavirus species but not Marburg virus and inhibits EBOV replication. Mechanistically, RETR1-2 recognizes EBOV-GP via its luminal domain, undergoes GP-induced oligomerization, and directs GP-containing ER membranes to lysosomes through canonical macro-autophagy. Using unbiased mass spectrometry, we identified TOLLIP as the key cytoplasm adaptor for RETR1-2, which also requires cooperation with the ER chaperone calnexin for EBOV-GP degradation. Notably, the PI3P-binding C2 domain of TOLLIP mediates its interaction with RETR1-2, and the EBOV-GP degradation occurs independently of ubiquitination, revealing an unexpected role for TOLLIP in ER-phagy. Furthermore, EBOV-VP40 antagonizes this pathway by selectively targeting RETR1-2 for macroautophagic degradation independently of TOLLIP, thereby restoring GP expression and viral infectivity. Nevertheless, RETR1-2 reciprocally degrades VP40 via a similar mechanism. Together, these findings define a calnexin–TOLLIP–RETR1-2 axis that links ER quality control to ER-phagy–mediated antiviral restriction and uncover a reciprocal host–virus arms race centered on selective macro-autophagy.

## Introduction

Ebola virus (EBOV) is a highly pathogenic filovirus that causes recurrent outbreaks of severe hemorrhagic fever with high mortality. Despite the availability of vaccines and antibody-based therapies, EBOV remains a persistent global health threat due to limited access to treatment, long-term viral persistence in immune-privileged sites, and the continual risk of re-emergence [1]. At the cellular level, productive EBOV infection critically depends on the synthesis, folding, and trafficking of its envelope glycoprotein (GP), a heavily glycosylated class I fusion protein that matures in the endoplasmic reticulum (ER) and is essential for viral entry and infectivity [2].

The ER is a central hub for protein quality control, and aberrant or misfolded viral glycoproteins are subject to host surveillance mechanisms. Selective autophagy of the endoplasmic reticulum (ER-phagy), mediated by dedicated ER-resident receptors, has emerged as an important pathway for maintaining ER proteostasis and restricting intracellular pathogens [3–6]. Notably, the RETREG1 (reticulophagy regulator 1) gene, also known as FAM134B, the first ER-phagy receptor identified, has been implicated in limiting EBOV infection, suggesting that EBOV-GP may be directly targeted by ER-phagy–dependent antiviral mechanisms [7]. However, how EBOV-GP is recognized by ER-phagy pathways, how the RETREG1 gene contributes to this process, and how ER quality control components interface with ER-phagy during viral infection remain poorly understood.

RETREG1/FAM134B belongs to the FAM134 family of ER-phagy receptors, which also includes RETREG2/FAM134A and RETREG3/FAM134C [8]. These receptors share an N-terminal reticulon homology domain (RHD) that drives ER membrane remodeling and a C-terminal LC3-interacting region (LIR) that links ER fragments to the autophagy machinery. The RETREG1 (RETR1) gene encodes two gene products, the full-length protein and an N-terminally truncated isoform 2, RETR1-2/FAM134B-2, which retains the C-terminal LIR but partially lacks the RHD. Although RETR1 and RETR1-2 are best known for their roles in starvation-induced macro-ER-phagy [9,10], RETR1 has been reported to exert both antiviral and proviral activities. The RETR1 gene has been reported to restrict pathogens such as EBOV [7], flaviviruses [11,12], porcine reproductive and respiratory syndrome virus (PRRSV) [13], and coronaviruses [14]. At the same time, this full-length protein in other contexts promotes viral replication by antagonizing host restriction factors [15,16]. The molecular basis underlying these opposing activities remains unresolved.

TOLLIP (Toll-interacting protein) is a phosphatidylinositol-3-phosphate (PI3P)–binding cytosolic autophagy adaptor that selectively targets aberrant membrane proteins for lysosomal degradation, classically in a ubiquitination-dependent manner [17]. TOLLIP contains an N-terminal TOM1-binding domain (TBD), a central C2 domain that mediates PI3P-dependent membrane association, an intrinsically disordered region (IDR) involved in cargo sensing, and a C-terminal coupling of ubiquitin to ER degradation (CUE) domain that binds ubiquitin. Unlike canonical ER-phagy receptors that localize to ER subdomains and engage autophagy-related 8 (ATG8)/LC3 family proteins via LIR motifs to engulf ER fragments into autophagosomes, TOLLIP has been primarily linked to endolysosomal trafficking pathways, and its role in ER-phagy–mediated antiviral restriction remains unclear.

Here, we identify RETR1-2, but not RETR1, as a selective ER-phagy receptor for EBOV-GP that restricts EBOV infection. We demonstrate that RETR1-2 recruits TOLLIP through its C2 domain and cooperates with the ER chaperone calnexin to direct GP to macro-autophagy–dependent lysosomal degradation. Strikingly, we further show that EBOV viral protein 40 kDa (VP40) antagonizes this pathway by selectively targeting RETR1-2 for macroautophagic degradation. Notably, RETR1-2 reciprocally degrades VP40 via a similar mechanism. Together, our findings uncover a calnexin–TOLLIP–RETREG1-2 axis that links ER quality control to ER-phagy–mediated antiviral defense and reveal an unexpected host–virus arms race centered on selective macro-autophagy.

## Results

### RETR1-2 broadly inhibits ebolavirus GP expression

The authentic EBOV replication has been reported to increase up to 100-fold in mice after knocking out the RETR1 gene, accompanied by strong upregulation of GP and VP40 protein expression in these mouse embryonic fibroblasts (MEFs) [7].

To identify the RETR1 gene product that inhibits the EBOV protein expression, we tested the activity of the full-length RETR1 and the truncated isoform RETR1-2. In addition, we tested the other known ER-phagy receptors, including testis-expressed protein 264 (TEX264), the reticulon 3 long isoform (RTN3L), Atlastin-3 (ATL3), cell cycle progression protein 1 (CCPG1), and secretory 62 (SEC62). When EBOV-GP was expressed with them in HEK293T cells, only RETR1-2 reduced the GP expression by 10-fold, as determined by Western blotting (WB) (**Fig. 1A**).

**Figure 1.**
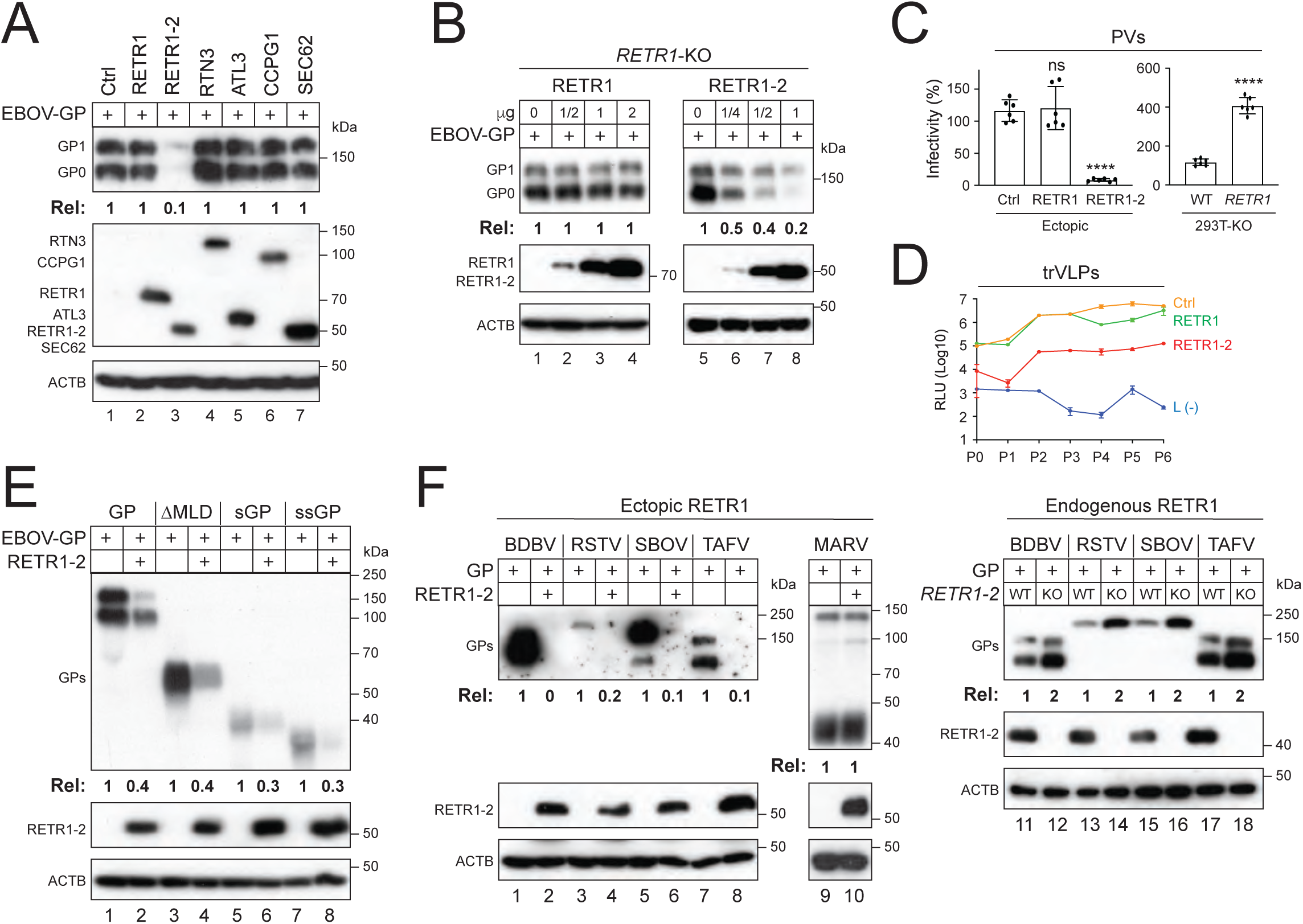
RETR1-2 broadly inhibits ebolavirus GP expression. **(A)** EBOV-GP was expressed with the indicated ER-phagy receptors in HEK293T cells, and protein expression was analyzed by WB. **(B)** EBOV-GP was expressed with full-length RETR1 or its truncated isoform RETR1-2 in HEK293T RETR1-KO cells, and protein expression was analyzed by WB. **(C)** HIV-1 firefly luciferase reporter pseudovirions (PVs) bearing EBOV-GP were produced in HEK293T WT cells expressing RETR1, RETR1-2, or a control vector, or in RETR1-KO cells. Viral infectivity was quantified following infection of HEK293T target cells by measuring intracellular luciferase activity. Data are presented as relative values, with infectivity of control PVs or WT cell-derived PVs set to 100. **(D)** EBOV replication- and transcription-competent virus-like particles (trVLPs) were generated (P0) and serially passaged six times (P1–P6) in HEK293T cells expressing ectopic RETR1, RETR1-2, or a control (empty vector). Viral replication was quantified by Renilla luciferase activity. trVLPs produced in the absence of EBOV-L [L(-)] served as a negative control. RLU, relative light unit. **(E)** EBOV-GP and its mucin-like domain deletion mutant (ΔMLD), soluble GP (sGP), and small soluble GP (ssGP) were expressed with RETR1-2 in HEK293T cells, and protein expression was analyzed by WB. **(F)** GPs from the indicated ebolavirus species and Marburg virus (MARV) were expressed with RETR1-2 in HEK293T cells or expressed in HEK293T WT and RETR1-KO cells. GP expression was analyzed by WB using anti-EBOV-GP antibodies. GP expression levels in panels (A), (B), (E) and (F) were quantified by ImageJ and presented as relative values. Error bars in (C) represent SEMs from three independent experiments (n = 3). *p < 0.05, **p < 0.01, ***p < 0.001, ****p < 0.0001; ns, not significant. All experiments were repeated at least three times; representative results are shown.

We then used a HEK293T RETR1-knockout (KO) cell line, where both RETR1 and RETR1-2 expression are disrupted [5]. When EBOV-GP was expressed with increasing amounts of RETR1 and RETR1-2, only RETR1-2 suppressed the GP expression in a dose-dependent manner (**Fig. 1B**, lanes 5-8).

We next evaluated the antiviral activity of RETR1-2 using infection-based assays. Human immunodeficiency virus type I (HIV-1) pseudovirions (PVs) bearing EBOV-GP produced from cells expressing RETR1-2 exhibited up to a 20-fold reduction in infectivity, whereas RETR1 had no effect (**Fig. 1C**, left). Conversely, PVs produced from the RETR1-KO cells displayed a 4-fold increased infectivity (**Fig. 1C**, right). Using Ebola virus replication-and transcription-competent virus-like particles (trVLPs), which recapitulate the full EBOV life cycle [18], RETR1-2 consistently suppressed viral replication by more than ten-fold over six consecutive passages, whereas RETR1 again showed no activity (**Fig. 1D**).

Among the other forms of EBOV-GP, RETR1-2 also reduced the expression of the soluble GP (sGP), small soluble GP (ssGP), and GP lacking the mucin-like domain (ΔMLD), indicating that its activity is independent of the MLD (**Fig. 1E**). RETR1-2 also suppressed GP expression from multiple ebolavirus species, including Sudan, Bundibugyo, Taï Forest, and Reston viruses, but not Marburg virus (**Fig. 1F**, lanes 1-10).

We then expressed these ebolavirus GPs in the RETR1-KO cells and found that their expression was strongly increased compared to wild-type (WT) cells (**Fig.1F**, lanes 11-18), confirming the RETR1-2 activity at endogenous levels.

Together, these data demonstrate that RETR1-2 broadly inhibits ebolavirus GP expression.

### RETR1-2 interacts with EBOV-GP via the luminal loop

RETR1 contains an N-terminal luminal loop, two clusters of transmembrane helices forming a reticulon homology domain (RHD), and a C-terminal LC3-interacting region (LIR) in the cytoplasm required for autophagosome recruitment (**Fig. 2A**, top; **Fig. S1A**). In contrast, RETR1-2 lacks the first cluster of transmembrane helices and therefore does not possess a functional RHD.

**Figure 2.**
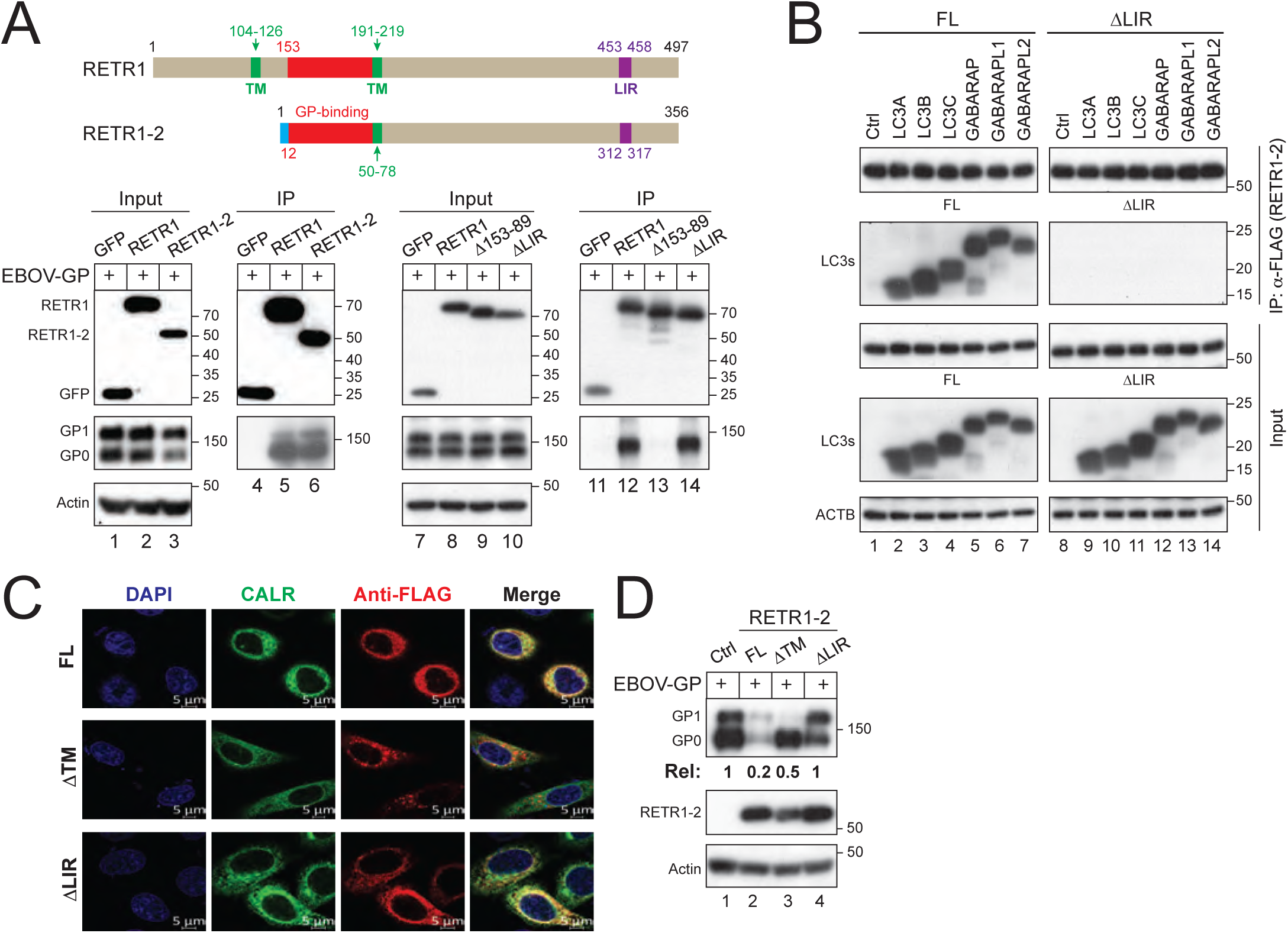
RETR1-2 interacts with EBOV-GP via the luminal loop. **(A)** Schematic representation of RETR1 and RETR1-2 showing transmembrane (TM) clusrer(s) (green), EBOV-GP binding domain (red), and LC3-interacting region (LIR; pink). Interaction of EBOV-GP with RETR1 or RETR1-2, and the deletion mutants of RETR1 (Δ153-89, ΔLIR) was analyzed by co-immunoprecipitation (co-IP) following expression in HEK293T cells; GFP served as a negative control. **(B)** FLAG-tagged RETR1-2 full-length (FL) and ΔLIR were co-expressed with HA-tagged LC3/GABARAP family proteins in HEK293T cells. Proteins were immunoprecipitated using anti-FLAG antibodies and analyzed by WB. **(C)** CALR-GFP was co-expressed with FLAG-tagged RETR1-2 FL, ΔLIR, or transmembrane-deleted mutant (ΔTM) in HeLa cells. Cells were stained with anti-FLAG antibodies followed by Alexa Fluor 647-conjugated secondary antibodies and analyzed by confocal microscopy (scale bar, 5 μm). **(D)** EBOV-GP was co-expressed with RETR1-2 FL, ΔLIR, or ΔTM in HEK293T cells, and protein expression was analyzed by WB. EBOV-GP expression levels in (D) were quantified by ImageJ. All experiments were repeated at least three times; representative results are shown.

Co-immunoprecipitation assays demonstrated that both RETR1 and RETR1-2 robustly interacted with EBOV-GP, whereas a control, green fluorescent protein (GFP), did not (**Fig. 2A**, lanes 4-6). We then deleted the shared luminal loop region in RETR1, residues 153-89, and found that this Δ153-89 mutant could no longer bind to EBOV-GP (**Fig. 2A**, lane 13). Thus, this luminal loop region (**Fig. 2A**, red; **Fig. S1B**) serves as the GP-binding domain that recruits GP for degradation from the ER.

To confirm that RETR1-2 functions as an ER-phagy receptor, we assessed the role of its LIR motif. RETR1-2, but not its LIR-deleted mutant (ΔLIR), interacted with LC3 and GABARAP family proteins (**Fig. 2B**), confirming that the LIR is required for autophagosome binding. Confocal microscopy showed that the full-length and ΔLIR proteins localized with the ER marker calreticulin (CALR), whereas the transmembrane-domain (TM) deletion mutant (ΔTM) failed to do so, confirming the ER localization of RETR1-2 via its TM (**Fig. 2C**).

Functionally, ΔLIR completely lost the ability to reduce EBOV-GP expression, while ΔTM exhibited a marked but incomplete loss of activity (**Fig. 2D**), indicating that both ER localization and LIR-mediated autophagosome recruitment are required.

These results demonstrate that RETR1-2 inhibits EBOV-GP expression via ER localization and LIR motif.

### RETR1-2 degrades EBOV-GP via canonical macro-autophagy

To understand how RETR1-2 inhibit EBOV-GP expression, we tried to block the RETR1-2 activity via pharmacological inhibition. Inhibiton of lysosomal or autophagic function (bafilomycin A1, concanamycin A, 3-MA, LY294002), but not proteasomal inhibition (lactacystin, MG132), abolished RETR1-2–mediated GP degradation (**Fig. 3A**). Consistently, EBOV-GP accumulated in LAMP1 (lysosome-associated membrane protein 1)-positive compartments upon RETR1-2 expression in the presence of lysosomal inhibitors (**Fig. 3B**).

**Figure 3.**
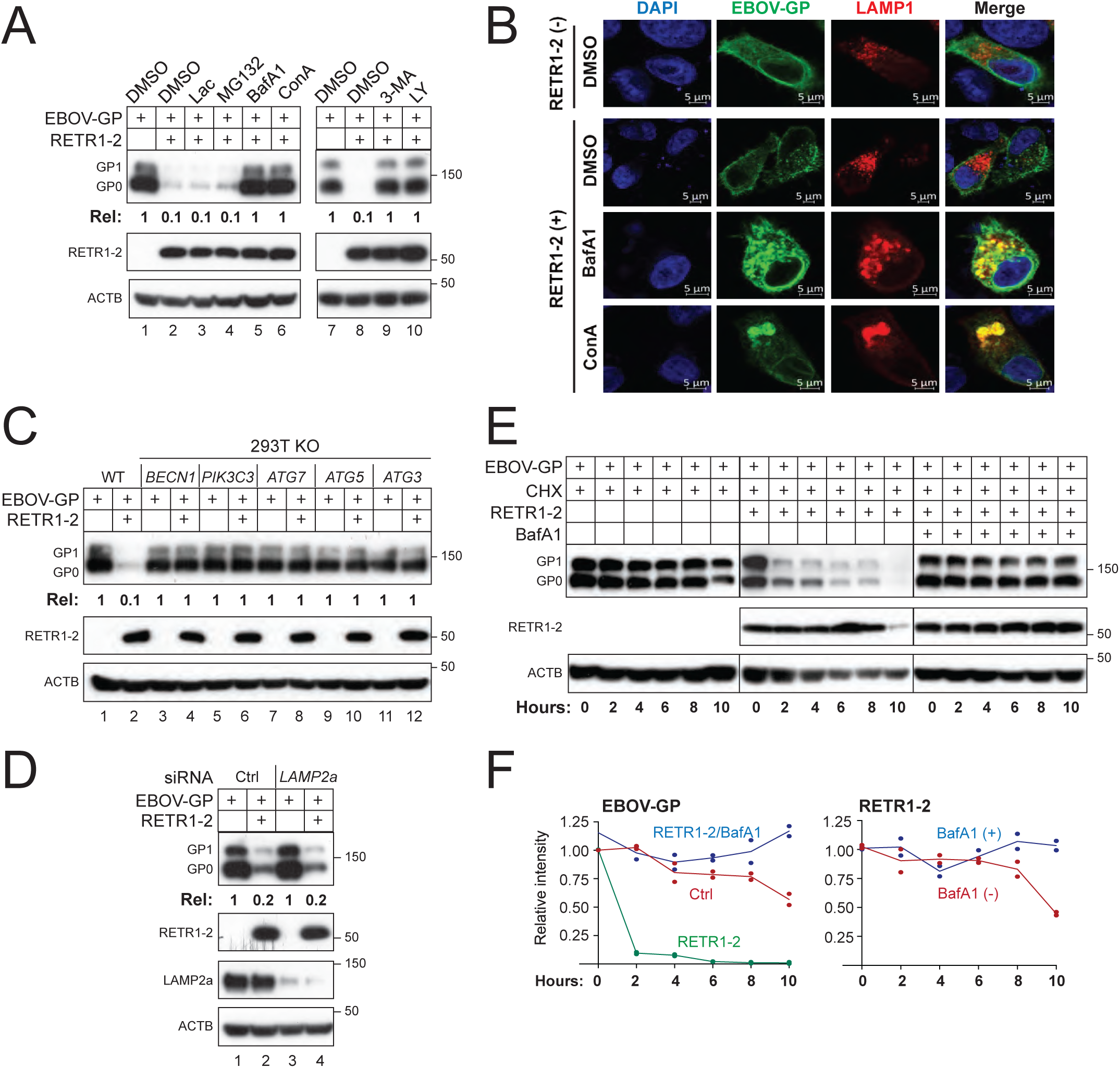
RETR1-2 mediates EBOV-GP degradation via canonical macro-autophagy. **(A)** EBOV-GP was expressed with RETR1-2 in HEK293T cells and treated with lactacystin (20 μM), MG132 (20 μM), bafilomycin A1 (BafA1; 100 nM), concanamycin A (ConA; 20 nM), 3-methyladenine (3-MA; 10 mM), or LY294002 (LY; 20 μM). Protein expression was analyzed by WB. **(B)** EBOV-GP-GFP and LAMP1-mCherry were co-expressed in HeLa cells in the absence of RETR1-2, or in the presence of RETR1-2 treated with ConA and BafA1. Colocalization was examined by confocal microscopy (scale bar, 5 μm). **(C)** EBOV-GP and RETR1-2 were expressed in HEK293T WT cells or cells lacking indicated key autophagy components, and protein expression was analyzed by WB. **(D)** EBOV-GP and RETR1-2 were expressed in HEK293T cells treated with LAMP2A-specific siRNA, and protein expression was analyzed by WB. **(E)** EBOV-GP was expressed with RETR1-2 in HEK293T cells, followed by cycloheximide (CHX; 50 μM) chase in the presence or absence of BafA1 (100 nM). Cells were harvested at the indicated time points and analyzed by WB. **(F)** EBOV-GP and RETR1-2 protein levels in (E) were quantified by ImageJ and expressed relative to time 0. EBOV-GP expression levels in (A), (C), and (D) were quantified by ImageJ. All experiments were repeated at least three times; representative results are shown.

To determine whether macro-autophagy is required for EBOV-GP degradation, we examined RETR1-2 activity in cells lacking key autophagy components. RETR1-2 failed to reduce EBOV-GP expression in cells deficient in PIK3C3/VPS34, BECN1, ATG3, ATG5, or ATG7 (**Fig. 3C**). In contrast, knockdown of LAMP2A did not impair RETR1-2 activity (**Fig. 3D**), excluding chaperone-mediated autophagy (CMA).

Cycloheximide chase assays revealed that EBOV-GP expressed alone exhibited a half-life of approximately 10 hours (h), whereas co-expression of RETR1-2 reduced the GP half-life to less than 2 h, an effect reversed by bafilomycin A1 (**Fig. 3E-F**). RETR1-2 itself remained stable throughout the chase period, indicating that it is not co-degraded.

Together, these data further establish RETR1-2 as a bona fide ER-phagy receptor that employs canonical macro-autophagy to degrade EBOV-GP.

### RETR1-2 recruits TOLLIP to mediate EBOV-GP degradation independently of ubiquitination

To identify cofactors involved in RETR1-2–mediated ER-phagy, we performed immunoprecipitation–mass spectrometry (IP-MS) analysis of RETR1-2 complexes. We identified 10 ER-resident proteins, including ER chaperones (CANX, PDIA3), ER-phagy receptors (RETR1, RETR3, Sec62), and others (SEC61A1, SEC61B, SEC63, CEPT1, RTN4) (**Fig. 4A**). The autophagic protein LC3B and late endosome protein RAB7A were also found in this complex, consistent with the role of RETR1-2 as an ER-phagy receptor. Notably, we identified the cytosolic autophagy adaptor TOLLIP, prompting further investigation.

**Figure 4.**
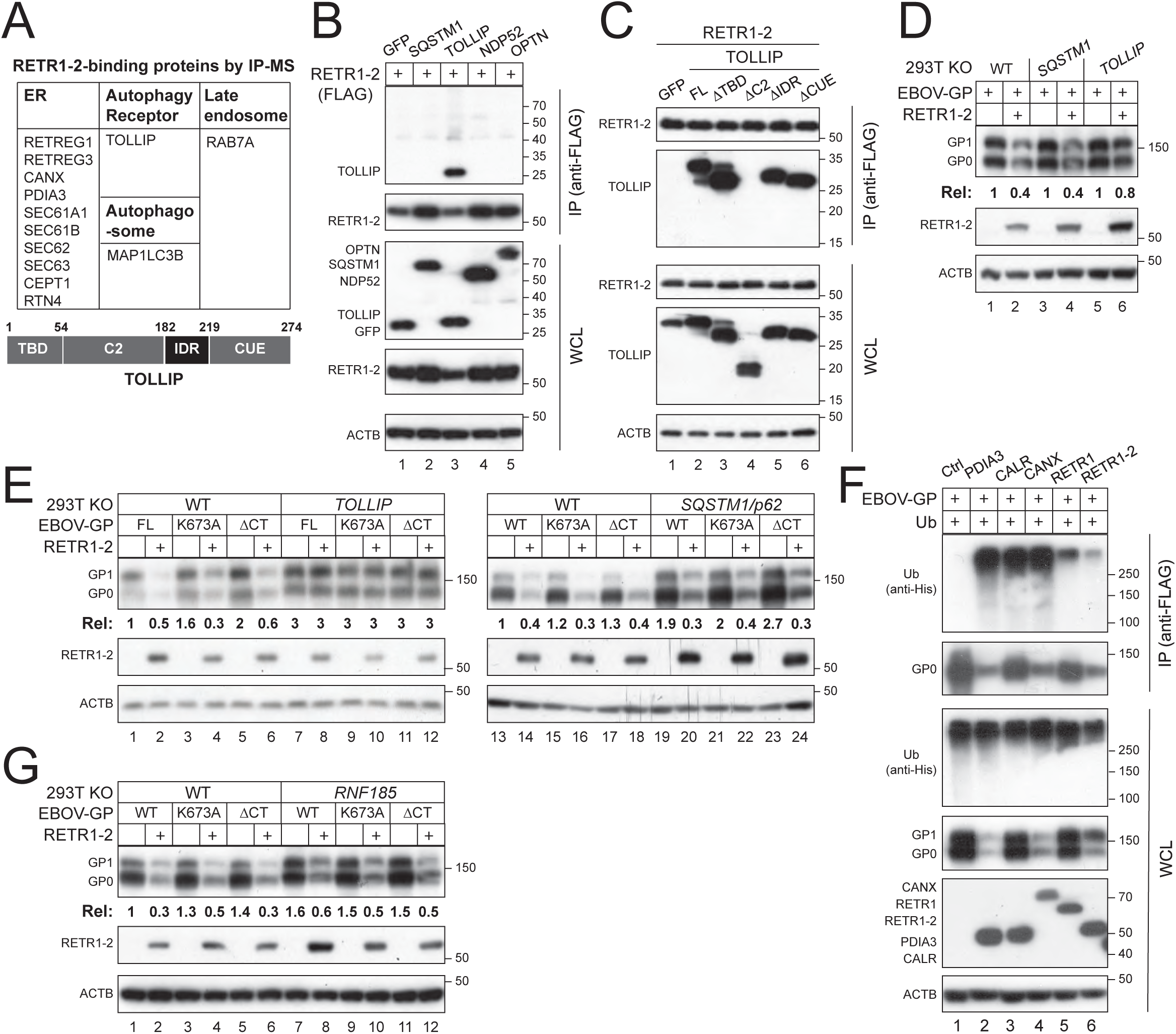
RETR1-2 recruits TOLLIP to mediate EBOV-GP degradation independently of ubiquitination. **(A)** RETR1-2–binding proteins identified by IP-MS are listed (top). A schematic of TOLLIP domain organization is shown (bottom), including the TOM1-binding domain (TBD), C2 domain, intrinsically disordered region (IDR), and CUE domain. **(B)** EBOV-GP and RETR1-2 were co-expressed with the indicated cytosolic autophagy receptors, and protein interactions were analyzed by co-IP. **(C)** RETR1-2 was co-expressed with TOLLIP full-length (FL) and indicated deletion mutants, and interactions were analyzed by co-IP. **(D)** EBOV-GP and RETR1-2 were expressed in HEK293T WT, SQSTM1-KO, or TOLLIP-KO cells, and protein expression was analyzed by WB. **(E)** EBOV-GP WT, K673A mutant, or cytoplasmic-tail deletion mutant (ΔCT) was expressed with RETR1-2 in WT, TOLLIP-KO, or SQSTM1-KO cells, followed by WB analysis. **(F)** EBOV-GP was expressed with the indicated ER chaperones, RETR1, and RETR1-2 in HEK293T cells, and GP polyubiquitination was analyzed by co-IP. **(G)** EBOV-GP was expressed with RETR1-2 in HEK293T WT or RNF185-KO cells, and protein expression was analyzed by WB. EBOV-GP expression levels in (D), (E), and (G) were quantified by ImageJ. All experiments were repeated at least three times; representative results are shown.

Co-immunoprecipitation assays confirmed a specific interaction between RETR1-2 and TOLLIP, whereas other autophagy adaptors (SQSTM1/p62, NDP52, OPTN) did not interact (**Fig. 4B**). Domain-mapping analyses were conducted by deleting the TBD, C2, IDR, or CUE region of TOLLIP and revealed that C2 deletion abolished its binding to RETR1-2, indicating that RETR1-2 engages TOLLIP through its C2 region (**Fig. 4C**).

Genetic ablation of TOLLIP, but not SQSTM1, eliminated RETR1-2–mediated EBOV-GP degradation (**Fig. S2**, **Fig. 4D**). Notably, RETR1-2 efficiently degraded EBOV-GP mutants lacking the ubiquitinated lysine (K673A) or the entire cytoplasmic tail (CT) where K673 locates, but only in TOLLIP-competent cells (**Fig. 4E**). Unlike ER chaperones PDIA3, CALR and CANX that could induce strong EBOV-GP polyubiquitination via RNF185 [6], both RETR1 and RETR1-2 exhibited little activity (**Fig. 4F**). Consistently, RETR1-2 degraded GP normally in the absence of its E3 ubiquitin ligase RNF185 (**Fig. 5G**).

**Figure 5.**
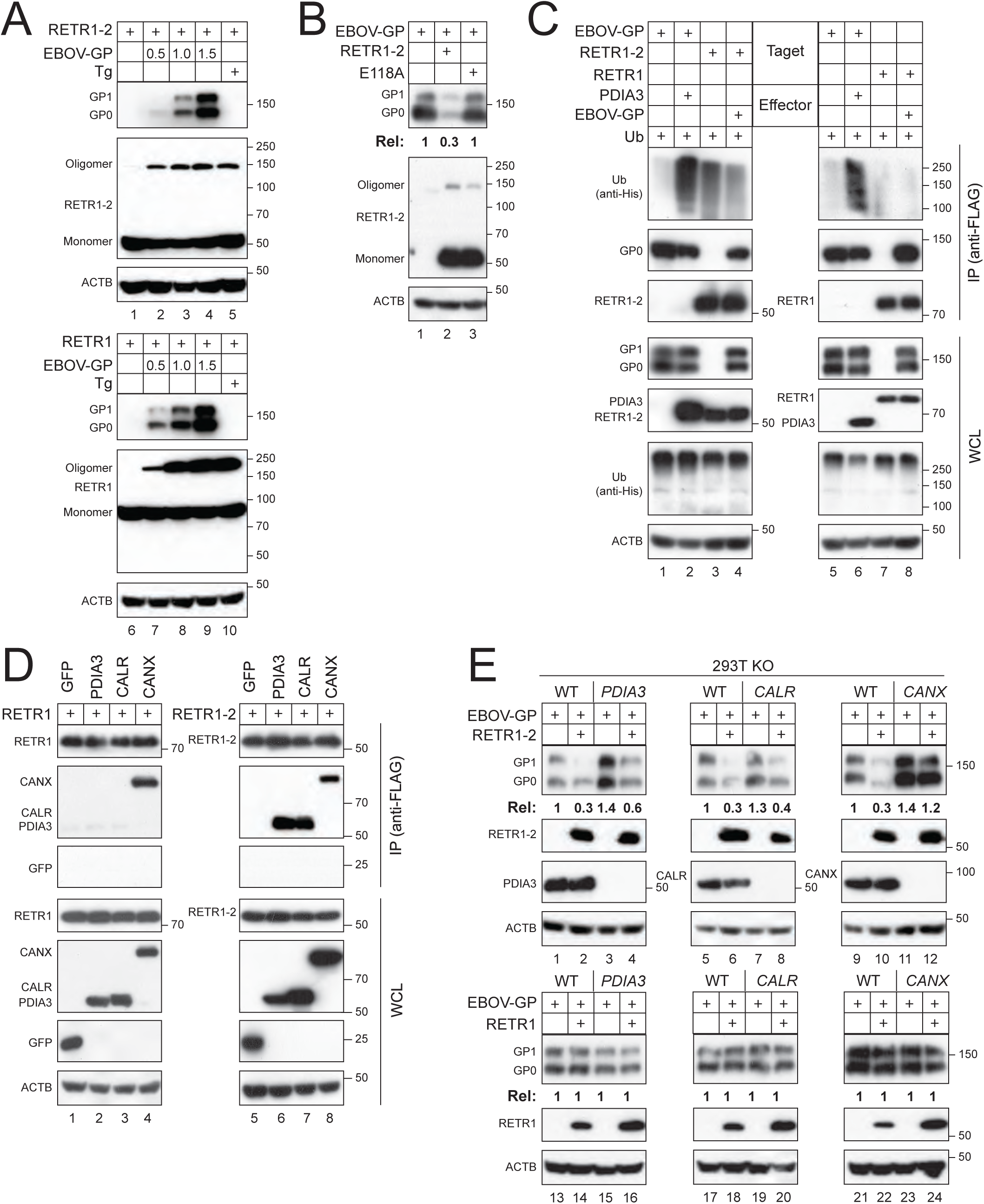
RETR1-2-mediated EBOV-GP degradation requires RETR1-2 oligomerization and CANX. **(A)** RETR1 or RETR1-2 was expressed with increasing amounts of EBOV-GP in HEK293T cells or expressed alone, followed by thapsigargin (Tg) treatment. RETR1 and RETR1-2 oligomerization was analyzed by WB. **(B)** EBOV-GP was expressed with RETR1-2 WT or mutant E118A. EBOV-GP expression and RETR1-2 oligomerization were analyzed by WB. **(C)** EBOV-GP was expressed with PDIA3, and alternatively, RETR1 and RETR1-2 were expressed with EBOV-GP in HEK293T cells. Polyubiquitination of EBOV-GP, RETR1, and RETR1-2 was analyzed by co-IP. **(D)** RETR1 and RETR1-2 were co-expressed with the indicated ER chaperones in HEK293T cells, and their interactions were analyzed by co-IP. **(E)** EBOV-GP was expressed with RETR1 or RETR1-2 in HEK293T WT and indicated knockout cells. Protein expression was determined by WB. EBOV-GP expression levels in (B) and (E) were quantified by ImageJ. All experiments were repeated at least three times; representative results are shown.

Together, these findings demonstrate that TOLLIP is essential for RETR1-2–mediated ER-phagy of EBOV-GP and that this process is independent of cargo ubiquitination.

### RETR1-2 oligomerization is required for EBOV-GP degradation

Because RETR1 oligomerization is required for ER-phagy and is regulated by cholesterol binding [19,20], we examined whether RETR1-2 oligomerization contributes to GP degradation. EBOV-GP expression induced oligomerization of both RETR1 and RETR1-2, like thapsigargin treatment (**Fig. 5A**).

Mutation of the conserved cholesterol-binding residue (E118A) markedly impaired RETR1-2 oligomerization and abolished its ability to degrade EBOV-GP (**Fig. 5B**). RETR1-2 itself was not polyubiquitinated during this process (**Fig. 5C**).

These results identify E118 as a critical determinant for RETR1 oligomerization and demonstrate that RETR1-2 relies on oligomerization—rather than ubiquitination—to initiate ER-phagy of EBOV-GP.

### RETR1-2 relies on calnexin to mediate EBOV-GP degradation

To confirm the RETR1-2 binding to ER chaperones identified from IP-MS (**Fig. 4A**), we expressed RETR1 and RETR1-2 with GFP, PDIA3, CALR, and CANX in HEK293T cells. Co-immunoprecipitation assays showed that RETR1-2 interacts with CANX, CALR, and PDIA3, whereas RETR1 interacts only with CANX (**Fig. 5D**).

To determine how ER chaperones contribute to RETR1-2–mediated EBOV-GP degradation, we co-expressed EBOV-GP with either RETR1 or RETR1-2 in HEK293T WT, PDIA3-KO, CALR-KO, or CANX-KO cells. As expected, RETR1 failed to reduce EBOV-GP expression in all cell lines tested (**Fig. 5E**, lanes 13–24). In contrast, RETR1-2 retained its ability to decrease EBOV-GP expression in PDIA3-KO and CALR-KO cells, comparable to WT cells, but this activity was completely abolished in CANX-KO cells (**Fig. 5E**, lanes 1–12).

Recently, we reported that EBOV-GP is targeted by the mannosidase alpha class 1B member 1 (MAN1B1)-Membralin/TMEM259 axis to ER-phagy [5]. However, RETR1-2 could still decrease EBOV-GP expression in MAN1B1-KO and Membralin-KO cells (**Fig. S3**).

These results demonstrate that RETR1-2–mediated EBOV-GP degradation specifically depends on CANX, but not on CALR, PDIA3, MAN1B1, or Membralin.

### EBOV antagonizes RETR1-2–mediated ER-phagy via VP40

To determine how EBOV counteracts RETR1-2, we assessed the major EBOV proteins for their ability to modulate RETR1-2 expression. EBOV encodes seven structural proteins: nucleoprotein (NP), VP35, VP40, GP, VP30, VP24, and the RNA-dependent RNA polymerase (L). Among these, only VP40 selectively reduced RETR1-2 protein levels in a dose-dependent manner, without affecting RETR1 (**Fig. 6A-B**).

**Figure 6.**
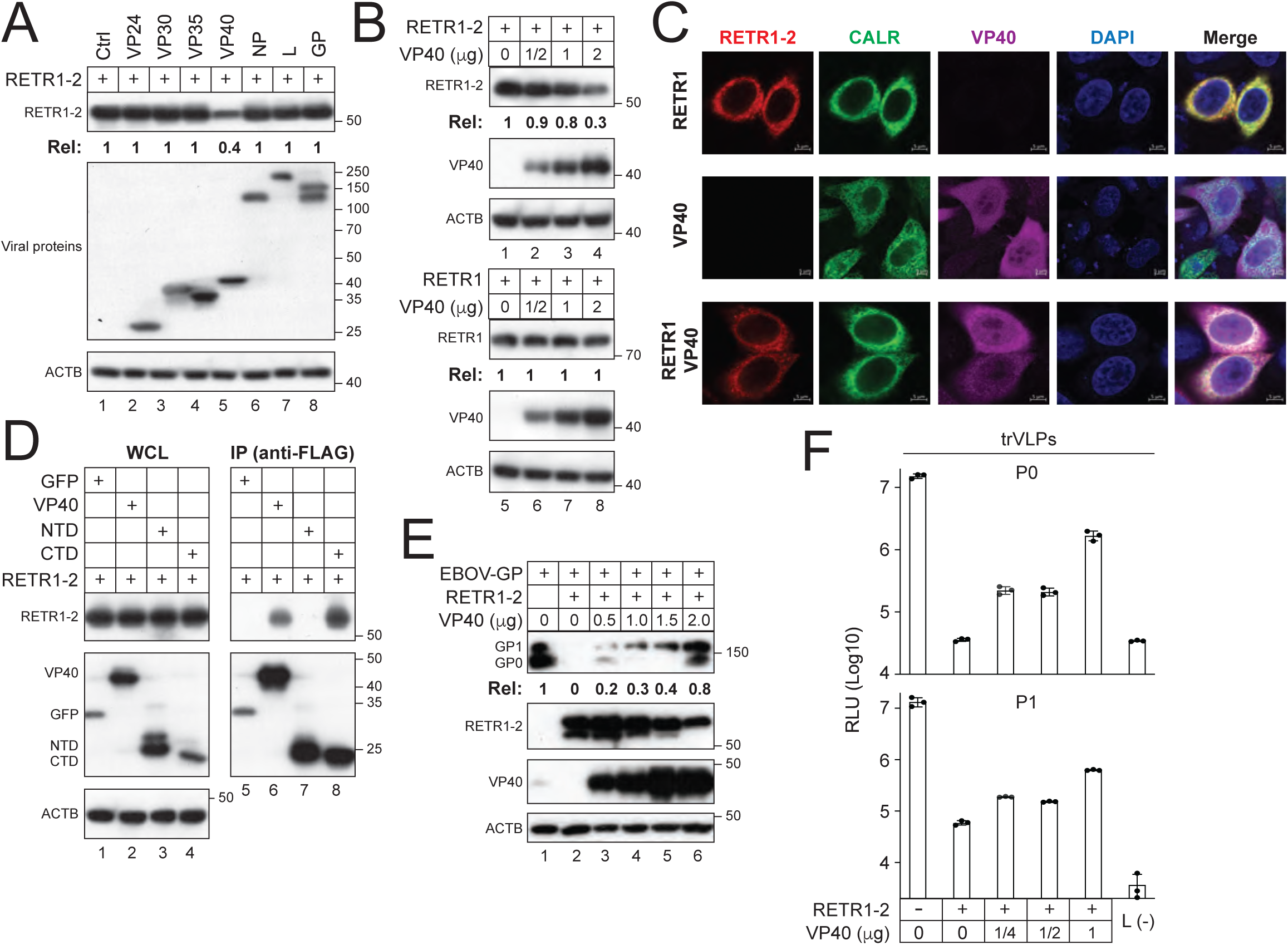
EBOV antagonizes RETR1-2 via VP40. **(A)** RETR1-2 was expressed with individual EBOV proteins in HEK293T cells, and protein expression was analyzed by WB. **(B)** RETR1 or RETR1-2 was expressed with increasing amounts of VP40, and protein expression was analyzed by WB. **(C)** RETR1-2 and VP40 were expressed alone or together in HeLa cells expressing CALR-GFP. Subcellular localization was examined by confocal microscopy (scale bar, 5 μm). **(D)** RETR1-2 was expressed with full-length VP40 or its N-terminal (NTD) or C-terminal (CTD) domain, and interactions were analyzed by co-IP. **(E)** EBOV-GP and RETR1-2 were co-expressed with increasing amounts of VP40 in HEK293T cells, and protein expression was analyzed by WB. **(F)** EBOV trVLPs were generated (P0) and passaged (P1) in HEK293T cells expressing ectopic RETR1-2 and increasing amounts of VP40. Viral replication was quantified as before. Protein expression levels in (A), (B), and (E) were quantified by ImageJ. All experiments were repeated at least three times; representative results are shown.

Confocal microscopy revealed that RETR1-2 recruits VP40 to the ER, where both proteins colocalized with the ER marker CALR (**Fig. 6C**). Domain-mapping experiments further demonstrated that RETR1-2 interacts with the C-terminal domain (CTD), but not the N-terminal domain (NTD), of VP40 (**Fig. 6D**). Functionally, VP40 restored the GP expression and viral infectivity in the presence of RETR1-2 in a dose-dependent manner (**Fig. 6E-F**). Together, these results indicate that VP40 antagonizes RETR1-2–mediated ER-phagy.

### VP40 targets RETR1-2 for macro-autophagic degradation

To define the mechanism by which VP40 counteracts RETR1-2, they were co-expressed in HEK293T cells and treated with proteasomal inhibitors (lactacystin, MG132) or lysosomal inhibitors (bafilomycin A1, concanamycin A). RETR1-2 expression was restored only by lysosomal inhibition, indicating that VP40 induces RETR1-2 degradation via the autophagy–lysosome pathway (**Fig. 7A**).

**Figure 7.**
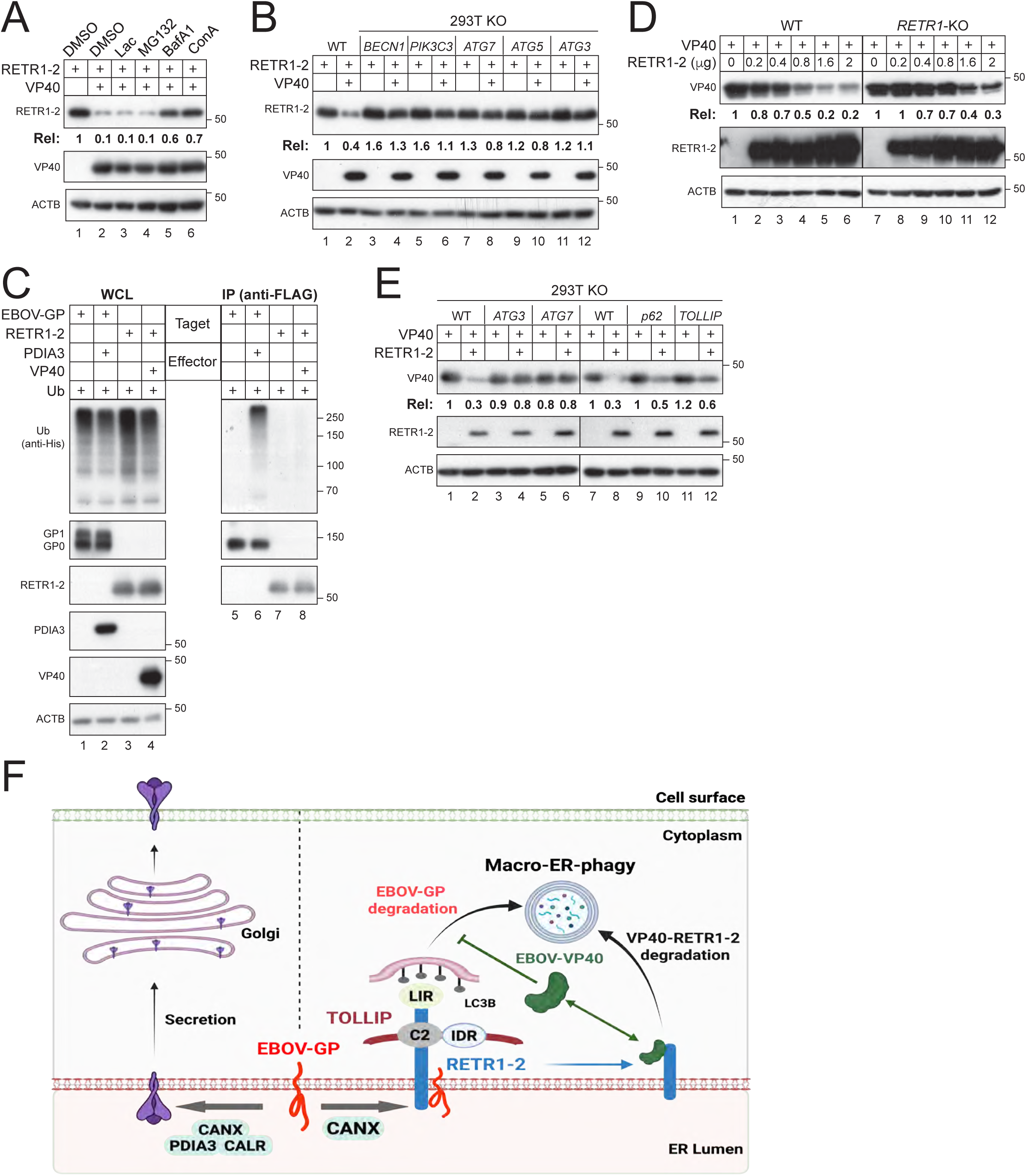
VP40 and RETR1-2 mutually target each other to macro-autophagy. **(A)** RETR1-2 and VP40 were co-expressed in HEK293T cells and treated with lactacystin, MG132, BafA1, or ConA, followed by WB analysis. **(B)** RETR1-2 and VP40 were expressed in HEK293T WT or indicated autophagy-deficient cell lines, and protein expression was analyzed by WB. **(C)** VP40 was expressed with RETR1 or RETR1-2, and EBOV-GP was expressed with PDIA3. Polyubiquitination was analyzed by co-IP. **(D)** VP40 was expressed with increasing amounts of RETR1-2 in HEK293T WT and RETR1-KO cells. Protein expression was analyzed by WB. **(E)** VP40 and RETR1-2 were expressed in HEK293T WT and indicated autophagy-deficient cell lines, and protein expression was analyzed by WB. **(F)** A model illustrating how RETR1-2 operates through two mechanistically distinct modes of action: a CANX–TOLLIP–dependent, ubiquitin-independent pathway for selective degradation of EBOV-GP, and a CANX–TOLLIP–independent pathway governing its own turnover with EBOV-VP40 during viral counteraction. Protein expression levels in (A), (B), (D), and (E) were quantified by ImageJ. All experiments were repeated at least three times; representative results are shown.

To further delineate this pathway, we examined VP40 activity in cells deficient in key autophagy components, including PIK3C3/VPS34, BECN1, ATG3, ATG5, and ATG7. In all cases, VP40 failed to reduce RETR1-2 expression, demonstrating that RETR1-2 degradation requires canonical macro-autophagy (**Fig. 7B**). Notably, VP40 did not induce RETR1-2 polyubiquitination, indicating that this process is ubiquitin-independent (**Fig. 7C**).

Consistent with prior observations that VP40 expression is elevated in RETR1-KO MEFs [7], we found that RETR1-2 also reduces VP40 expression in a dose-dependent manner in both WT and RETR1-KO HEK293T cells (**Fig. 7D**). This antiviral activity was abolished in ATG3-KO and ATG7-KO cells but remained intact in SQSTM1/p62-KO or TOLLIP-KO cells, indicating that this VP40 degradation also proceeds via macro-autophagy independently of SQSTM1/p62 and TOLLIP (**Fig. 7E**).

Collectively, these findings demonstrate that RETR1-2 and VP40 mutually target each other for macro-autophagic degradation. In contrast to RETR1-2–mediated GP degradation, this reciprocal degradation does not require TOLLIP.

## Discussion

A key insight from this study is that RETR1-2, but not full-length RETR1, acts as an ER-phagy receptor of EBOV-GP, despite both isoforms retaining the GP-binding region. Notably, EBOV-GP induces oligomerization of both RETR1 than RETR1-2, arguing against defective cargo engagement as the basis for RETR1 inactivity. Instead, the critical distinction appears to lie downstream of cargo recognition, at the level of coupling to the autophagy machinery. Recent work has shown that the RHD of FAM134 family proteins drives membrane curvature and oligomerization but can also constrain receptor dynamics and cargo processing [21]. Consistent with this, RETR1 may be optimized for curvature-driven, homeostatic ER turnover, whereas RETR1-2, lacking part of the RHD, is better suited to couple cargo recognition to selective autophagic degradation. This distinction becomes particularly relevant for cargos that are not efficiently polyubiquitinated. In this context, RETR1 fails to promote productive degradation, whereas RETR1-2 uniquely engages the adaptor TOLLIP to direct EBOV-GP to macro-autophagy in a ubiquitin-independent manner, enabling efficient clearance of “ubiquitin-silent” viral glycoproteins.

A central component of this pathway is the ER chaperone CANX, which appears to act upstream of RETR1-2 in substrate selection. CANX is a core ER quality-control factor that binds glycoproteins through their N-linked glycans and monitors their folding status as part of the calnexin/calreticulin cycle [22]. Importantly, recent studies have established a functional link between RETR1 and CANX in ER-phagy, demonstrating that CANX can act as a luminal cofactor that delivers misfolded or aggregation-prone glycoproteins to RETR1-dependent ER-phagy pathways [23–25]. Our data extend this model to antiviral defense by showing that RETR1-2–mediated EBOV-GP degradation strictly depends on CANX, but not on other ER chaperones such as CALR or PDIA3, suggesting a selective handoff mechanism. In this model, CANX recognizes EBOV-GP as a non-native or persistent client and recruits it into a RETR1-2–containing complex, thereby linking ER quality control to ER-phagy (**Fig. 7F**). This step likely provides substrate specificity, ensuring that viral glycoproteins are preferentially targeted while avoiding indiscriminate ER turnover. The requirement for CANX further supports a model in which luminal glycoprotein surveillance, rather than ubiquitination, serves as the primary trigger for this pathway.

Consistent with this, RETR1-2–mediated ER-phagy is TOLLIP-dependent but ubiquitin-independent, revealing a noncanonical mechanism of selective autophagy. TOLLIP, classically viewed as a ubiquitin-binding adaptor, here functions through its PI3P-binding C2 domain to couple RETR1-2–bound cargo to autophagic membranes. Thus, RETR1-2 integrates luminal cargo sensing via CANX with cytosolic membrane targeting via TOLLIP, establishing a bipartite recognition system that bypasses the need for ubiquitin signals. Importantly, RETR1-2 itself is not co-degraded, indicating that it acts catalytically to deliver cargo to autophagosomes.

In contrast, the reciprocal targeting of RETR1-2 and VP40 reveals a mechanistically distinct mode of macroautophagic degradation that does not require CANX, TOLLIP, or ubiquitination (**Fig. 7F**). This suggests that, unlike EBOV-GP, the RETR1-2–VP40 complex is recognized as an aberrant assembly capable of directly engaging the core autophagy machinery. One possibility is that VP40 binding alters RETR1-2 conformation or oligomeric state, converting it from a functional receptor into a degradable substrate. Alternatively, their interaction may generate higher-order complexes that are intrinsically targeted by ubiquitin-independent autophagy pathways. Notably, RETR1-2 and VP40 mutually target each other through distinct mechanisms: a CANX–TOLLIP–dependent pathway for viral glycoprotein clearance and a CANX–TOLLIP–independent pathway for reciprocal degradation. This bidirectional antagonism highlights selective macro-autophagy as a dynamic interface of host–virus conflict.

Consistent with this model, RETR1-2 interacts with the C-terminal domain of VP40, a region known to mediate membrane binding, oligomerization, and host factor interactions. The VP40 C-terminal domain contains basic patches that preferentially associate with negatively charged phospholipids, suggesting that RETR1-2 may recruit VP40 to specialized ER subdomains. This is supported by their ER colocalization and suggests that RETR1-2 functions not only as a cargo receptor but also as a spatial organizer that captures viral proteins at the ER.

Together, these findings support a model in which RETR1-2 operates through two mechanistically distinct modes of action: a CANX–TOLLIP–dependent, ubiquitin-independent pathway for selective degradation of viral glycoproteins, and a CANX–TOLLIP–independent pathway governing its own turnover with VP40 during viral counteraction. This dual functionality underscores the versatility of ER-phagy receptors and reveals how cargo identity dictates pathway selection, uncovering a bidirectional host–virus arms race centered on selective macro-autophagy.

## Experimental Section

### Antibodies and inhibitors

Commercial reagents include: MG132, Lactacystin (Lac), Concanamycin A (ConA), 3-Methyladenine (3-MA), Thapsigargin (TGN) and Cycloheximide (CHX) (Sigma-Aldrich, C2211, L6785, C9705, M9281, T9093, 5087390001); Bafilomycin A1 (BafA1, Santa Cruz Biotechnology, sc-201550); rabbit polyclonal anti-EBOV-GP and anti-EBOV-VP40 (Sino Biological, 40442-T48, 40446-T48); mouse monoclonal anti-Myc (CST, 2276S); mouse monoclonal anti-FLAG and anti-HA (Sigma-Aldrich, F3165, H3663); mouse monoclonal anti-ACTB/β-actin (CST, 3700S); horseradish peroxidase (HRP)-conjugated goat anti-mouse IgG and anti-rabbit IgG (Jackson ImmunoResearch Laboratories, 115-035-003, 111-035-003).

### Cell lines

Human embryonic kidney (HEK) 293 cell line transformed with SV40 large T antigen (293T), human cervical carcinoma cell line HeLa and African green monkey kidney epithelial cell line Vero E6 were purchased (ATCC, CRL-3216, CRM-CCL-2, CRL-1586). The exact purchase dates of these cell lines could not be verified, but this does not affect the conclusions. All cell lines were confirmed to be free of contamination.

All these cells were maintained in Dulbecco’s modified Eagle medium (DMEM; Thermo Fisher Scientific, 11965092) supplemented with 10% fetal bovine serum (FBS) and 1% penicillin-streptomycin (pen-strep; Thermo Fisher Scientific, 10378016) and cultivated at 37°C in the humidified atmosphere in a 5% CO_2_ incubator.

### CRISPR-Cas9 knockout (KO) cell lines

The HEK293T MAN1B1-KO cell line was reported [26]. The HEK293T calnexin-KO, calreticulin-KO, and PDIA3-KO cell lines were reported [4]. The HEK293T RETREG1-KO and membralin-KO cell lines were reported [5]. The HEK293T TOLLIP-KO cell line was reported [27]. The HEK293T RNF185-KO, SQSTM1/p62-KO, BECN1-KO, PIK3C3-KO, ATG7-KO, ATG5-KO, and ATG3-KO cell lines were reported [6,15].

### Plasmids

The pcDNA3.1-HiBiT-EBOV-GP, pcDNA3.1-HiBiT-EBOV-GPΔMLD, pcDNA3.1-HiBiT-EBOV-sGP, pcDNA3.1-HiBiT-EBOV-ssGP, pcDNA3.1-Flag-EBOV-GP, pcDNA3.1-Flag-SEBOV-GP, pcDNA3.1-Flag-RETV-GP, pcDNA3.1-Flag-TAFV-GP, pcDNA3.1-Flag-BDBV-GP, pCAGGS-PDIA3-Myc, pCMV6-CALR-Myc, pCMV6-CANX-Myc expression vectors were reported previously [4,6]. pCDNA3.1-RETREG1-FLAG, pCMV6-RETREG1-2-Myc-Flag, pCMV6-LC3A-HA, pCMV6-LC3B-HA, pCMV6-LC3C-HA, pCMV6-GABARAP-HA, pCMV6-GABARAPL1-HA, and pCMV6-GABARAPL2-HA expression vectors were reported previously [5]. pCMV6-SQSTM1-HA, pCMV6-TOLLIP-HA, pCMV6-NDP52-HA, and pCMV6-OPTN-HA expression vectors were reported previously [27]. TOLLIP mutants were created by overlapping PCR. pCMV6-RTN3-Myc-Flag, pCMV6-ATL3-Myc-Flag, pCMV6-CCPG1-Myc-Flag, pCMV6-SEC62-Myc-Flag were ordered from Comatebio (CoME). RETREG1-2 mutants were created by PCR and *ASiSI*/*MluI* digestion followed by homologous recombination. pCAGGS-VP24-3 x Flag, pCAGGS-VP30-3 x Flag, pCAGGS-VP35-3 x Flag, pCAGGS-VP40-3 x Flag, pCAGGS-NP-3 x Flag, pCAGGS-L-3 x Flag were created by PCR and homologous recombination. All vectors constructed by us were confirmed by Sanger DNA sequencing. Detailed experimental procedures and primer sequences for the construction of these vectors are available upon request. Plasmids were prepared using Maxiprep kits (TIANGEN Biotech, DP117).

### Transfection

HEK293T cells were cultured in 6-well plates and transfected with polyethyleneimine (PEI; Polysciences, 23966-2). HeLa cells were transfected using Lipofectamine 3000 according to the manufacturer’s protocol (Thermo Fisher Scientific, L3000015). The total indicated plasmids were diluted into 200μL serum-free Opti-MEM (Thermo Fisher Scientific, 31985062) and mixed with transfection reagents. After 20 min of incubation at room temperature, these transfection complexes were added directly into the supernatant of each well. Media were replaced after 6 h and cell lysate was collected at 48 h unless otherwise noted.

### FLuc-psedovirion (PV) infection

EBOV-GP pseudotyped HIV-1 virions were produced from HEK293T WT and *RETREG2*-KO cells as we reported previously [28]. These cells were transfected with pNL-Luc-ΔEnv and a S protein expression vector in the presence or absence of RETREG1 or RETREG1-2 expression. After 48 h, viruses were collected from the culture supernatants and clarified by low-speed centrifugation. After being normalized by p24^Gag^ ELISA [29], an equal amount of viruses were used to infect Vero E6 cells. After 48 h of infection, cells were lysed, and viral infectivity was determined by measuring the intracellular luciferase activity using a firefly luciferase assay kit (US Everbright Inc, Cat# F6024).

### Production of EBOV trVLPs and infection

Experiments were performed as reported previously [18]. Briefly, to produce p0 trVLPs, HEK293T cells were seeded in 6-well plates at 4 x 10^5^ per well in 2 mL medium and cultured for 24 h. EBOV trVLP system plasmids were diluted from stocks into 125 μL Opti-MEM per well that included 125 ng pCAGGS-NP, 125 ng pCAGGS-VP35, 75 ng pCAGGS-VP30, 1,000 ng pCAGGS-L, 250 ng p4cis-vRNA-Rluc, and 250 ng pCAGGS-T7. The plasmids were combined with 7.5 μL TransIT^®^-LT1 Transfection Reagent (Mirus Bio, 2305). After incubation at room temperature for 15 min to promote complex formation, the transfection mixes were gently added into cell culture. After 24 h of transfection, the media was exchanged for 4 mL growth media with 5% FBS, and further incubated for 72 h. All supernatants from p0 producer cells were collected followed by centrifugation at 3,000 x g for 5 min and stored at −80 °C until use. To generate p1 target cells, HEK293T cells were transfected similarly with EBOV trVLP system plasmids but had p4cis-vRNA-Rluc and pCAGGS-T7 replaced with 250 ng pCAGGS-Tim1. Transfected p1 cells were infected with p0 trVLPs, and after 72 h, p1 trVLPs were collected and stored. P2 target cells were prepared similarly and infected with p1 trVLPs. Cells from p0, p1, and p2 were lysed to quantify virus production by Renilla-Glo^®^ Luciferase Assay System (Promega, E2820).

### Western blotting (WB)

Transfected cells were lysed in NP-40 (Beyotime, P0013F) at 4°C. After centrifugation at 12,000 × g for 10 min at 4°C, cytosolic fractions were collected and boiled with loading buffer (Solarbio Life Sciences, P1015). Proteins were separated by PAGE Gel FAST Preparation Kit (Epizyme, PG112) and transferred onto PVDF membranes. These membranes were blocked with 5% nonfat milk powder in TBST (Tris-buffered saline [20 mM Tris, pH 7.4,150 mM NaCl] containing 0.1% Tween 20; Solarbio Life Sciences, T8220) for 1 h at room temperature and probed by primary antibodies followed by HRP-conjugated secondary antibodies. The blotted proteins were detected by SuperSignal substrate (Thermo Fisher Scientific, 34580) using X-ray films (FUJI) as reported [30]. Films were scanned, and protein bands were quantified by ImageJ. Adobe Photoshop and Adobe Illustrator were used to generate the figures.

### Immunoprecipitation

After transfection of HEK293T cells cultured in a 6-cm dish, cells were lysed in 0.8 mL NP40 lysis buffer for 30 min on ice as we did previously [31]. After the removal of nuclei via low-speed centrifugation and collecting 100 μL as input, the remaining 700 μL lysate was incubated with anti-FLAG M2 Magnetic beads (Sigma-Aldrich, M8823) and rotated at 4 °C overnight. After being washed 3 times with 1 mL pre-cooled NP40 lysis buffer, proteins were removed from beads after boiling in 40 μL NP40 lysis buffer plus 15 μL sample loading buffer (5×) and analyzed by WB.

### In vivo polyubiquitination assay

HEK293T cells were seeded in 6-cm dishes and transfected with vectors expressing EBOV-GP and RETREG1 or RETREG1-2, PDIA3, CALR, CANX, in the presence of ubiquitin expression vector as we did previously [32]. After 48 h, cells in each dish were lysed in 600 μL NP40 buffer at 4°C for 30 min. After the removal of nuclei by low-speed centrifugation, 100 μL was collected as input, and the remaining 500 μL was incubated with anti-FLAG beads (Sigma-Aldrich, M8823) at 4°C overnight. Beads were washed three times with phosphate-buffered saline (PBS), then boiled in SDS-PAGE loading buffer and analyzed by WB.

### HiBiT blotting

Proteins were similarly separated by SDS-PAGE and transferred to a PVDF membrane. After that, proteins with HiBiT-tag were detected by Nano-Glo^®^ HiBiT Blotting System (Promega, N2410). Briefly, the membrane was incubated with LgBiT/buffer solution (Promega, N2421) for 2 h at room temperature with gentle rocking and then at 4 °C overnight. The membrane was incubated with 20 μL of substrate for 5 min at room temperature and placed between transparent plastic sheets for imaging.

### Mass spectrometry

HEK293T cells were transfected with either pFlag-FAM134B-2 or pFlag-Vector. Samples were collected at 36 hours post-transfection. Co-immunoprecipitation (Co-IP) was performed using anti-FLAG beads, and the immunoprecipitated samples were sent to the BGI TECH SOLUTIONS (BEIJING LIUHE) CO., LIMITED, for Nano LC-MS/MS and database search analysis. The raw data can be downloaded from iProX (project ID: IPX0015521000).

### Confocal microscopy

HeLa cells were seeded on a glass-bottom cell culture dish and transfected with various vectors using Lipofectamine 3000, as we did previously[33]. After 30 h, cells were fixed with 4% paraformaldehyde for 10 min, permeabilized with 0.1% Triton X-100 (Solarbio Life Sciences, T8200) for 10 min at room temperature, and then blocked with 5% bovine serum albumin (BSA; APPLYGEN, P1622) solution overnight at 4°C. Nuclei were stained with 4′,6-diamidino-2-phenylindole (DAPI) for 3-5 min and washed with PBS, and cells were observed and imaged under a confocal microscope (LSM880, Zeiss, White Plains, New York).

### Statistical analysis

All experiments were performed independently at least three times, with representative experiment being shown. GraphPad Prism (Graph Pad Software Inc., San Diego, CA, USA) was used for the data analysis. Data were presented as means ± standard error of measurements (SEMs) and represented by error bars. The significance of differences between samples was assessed by an unpaired two-tailed Student’s t test. A *p* value < 0.05 (*p* < 0.05) was statistically significant (*p < 0.05, \*\**p* < 0.01, \*\*\**p* < 0.001), and *p* > 0.05 was not significant (ns).

## Acknowledgments

We thank Feng Zhang for the reagent. We sincerely acknowledge members in the laboratory for their technical assistance and helpful advice.

## Data availability

All data supporting the findings of this study is available from the corresponding authors on reasonable request.

## Author Contributions

J.Z. and T.W. conducted most of the experiments with support from J.W., J.L.., and S.L. J.Z., T.W., and Y.H.Z. designed experiments. Y.H.Z. wrote the manuscript with input from all authors.

## Abbreviations

ATL3: Atlastin-3
ATG: autophagy-related
ATG3: autophagy-related protein 3
ATG5: autophagy-related protein 5
ATG7: autophagy-related protein 7
ATG8: autophagy-related protein 8
BafA1: bafilomycin A1
BECN1: beclin 1
CALR: calreticulin
CANX: calnexin CCPG1 cell cycle progression protein 1
CHX: cycloheximide
CMA: chaperone-mediated autophagy
ConA: concanamycin A
CT: cytoplasmic tail
CTD: C-terminal domain
CUE: coupling of ubiquitin to ER degradation
EBOV: Ebola virus
EBOV-GP: Ebola virus glycoprotein
ER: endoplasmic reticulum
ERAD: endoplasmic reticulum–associated degradation
ER-phagy: selective autophagy of the endoplasmic reticulum
FL: full-length
FAM134B: family with sequence similarity 134 member B
FAM134B-2: family with sequence similarity 134 member B isoform 2
GP: glycoprotein
GFP: green fluorescent protein
HIV-1: human immunodeficiency virus type 1
IDR: intrinsically disordered region
IP: immunoprecipitation
IP-MS: immunoprecipitation–mass spectrometry
KO: knockout
L: L-polymerase
LAMP1: lysosome-associated membrane protein 1
LAMP2A: lysosome-associated membrane protein 2A
LC3: microtubule-associated protein 1 light chain 3
LIR: LC3-interacting region
LY294002: phosphoinositide 3-kinase inhibitor
MAN1B1: mannosidase alpha class 1B member 1
MEFs: mouse embryonic fibroblasts
MLD: mucin-like domain
NP: nucleoprotein
NTD: N-terminal domain
OPTN: optineurin
PDIA3: protein disulfide isomerase family A member 3
PI3P: phosphatidylinositol-3-phosphate
PIK3C3: phosphatidylinositol 3-kinase catalytic subunit type 3
PRRSV: porcine reproductive and respiratory syndrome virus
PVs: pseudovirions
RETR1: RETREG1
RETR1-2: RETREG1-2
RETREG1: reticulophagy regulator 1
RETREG1-2: reticulophagy regulator 1 isoform 2
RHD: reticulon homology domain
RNF185: ring finger protein 185
RLU: relative light unit
RTN3L: reticulon 3 long isoform
SEC62: secretory 62
sGP: soluble glycoprotein
SQSTM1: sequestosome 1
ssGP: small soluble glycoprotein
TBD: TOM1-binding domain
TEX264: testis-expressed protein 264
Tg: thapsigargin
TM: transmembrane domain
TOLLIP: Toll-interacting protein
trVLPs: transcription- and replication-competent virus-like particles
VP40: EBOV viral protein 40 kDa
WT: wild type
ΔCT: cytoplasmic-tail deletion mutant
ΔLIR: LC3-interacting region deletion mutant
ΔMLD: mucin-like domain deletion mutant
ΔTM: transmembrane domain deletion mutant.

## Figure Legends

**Figure S1.**
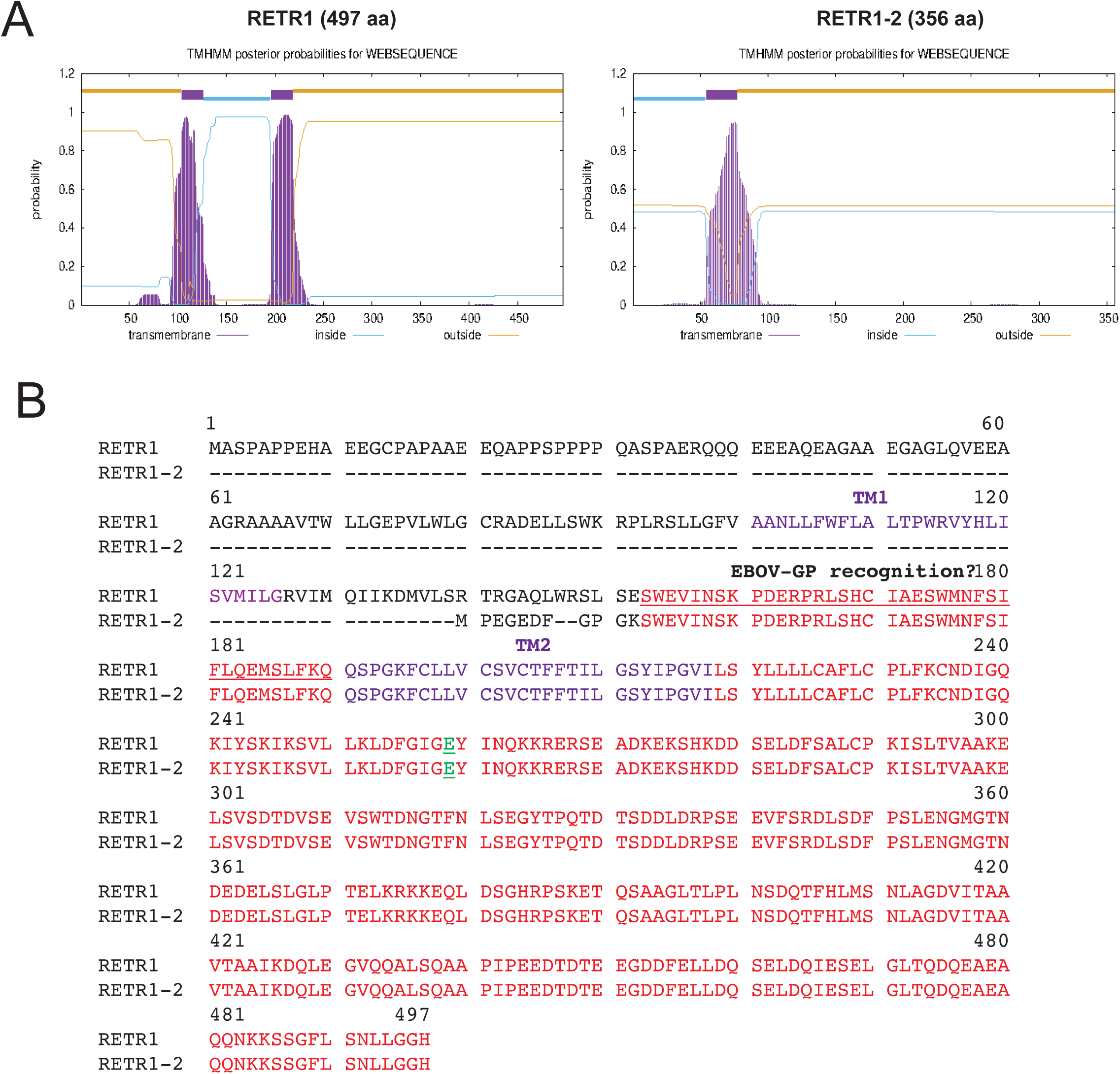
**(A)** The transmembrane helices of RETR1 and RETR1-2 were precited by TMHMM - 2.0 online tool. Their transmembrane domain, cytoplasmic domain (inside), and ER lumianl domain (inside) are shown. **(B)** The amino acid sequences of RETR1 and RETR1-2 are aligned. Their TM regions and the putattive EBOV-GP recognition seqence are inidicatd. The the key cholesterol-binding site that determines oligomerization (E259 in RETR1, or E118 in RETR1-2) is shown in green and underlined.

**Figure S2.**
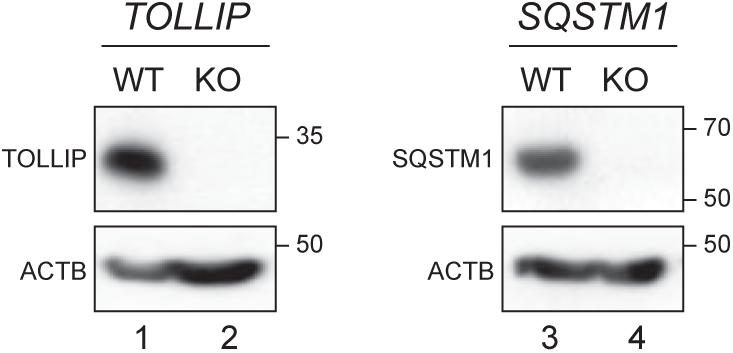
Generation of HEK293T TOLLIP-KO and SQSTM1-KO cells by CRISPR/Cas9.

**Figure S3.**
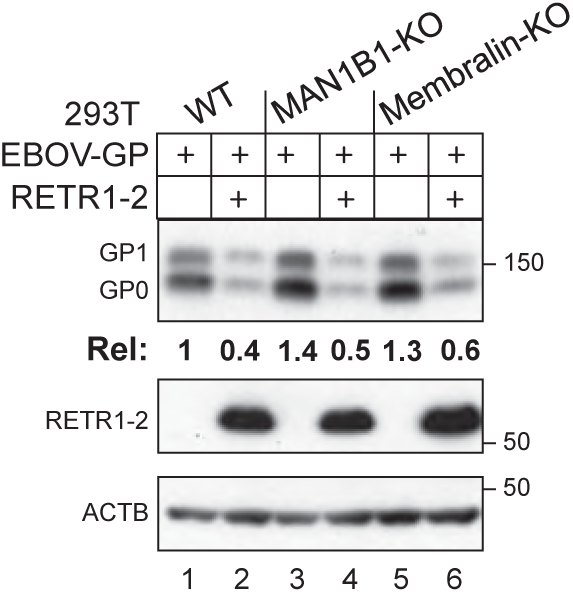
EBOV-GP and RETR1-2 were expressed in HEK293T and indicated KO cells and their expression was determined by WB.

